# Epigenetic memory astrocytes are likely an artifact of immune cell contamination

**DOI:** 10.1101/2025.04.04.647246

**Authors:** Michael R. O’Dea, Shane A. Liddelow

**Affiliations:** Institute for Translational Neuroscience, NYU Grossman School of Medicine, New York, NY., 10016, USA; Department of Neuroscience, NYU Grossman School of Medicine, New York, NY., 10016, USA; Department of Ophthalmology, NYU Grossman School of Medicine, New York, NY., 10016, USA; Parekh Center for Interdisciplinary Neurology, NYU Grossman School of Medicine, New York, NY., 10016, USA

## Abstract

Innate immune memory, in which prior immune stimuli can “train” certain immune cells to respond more aggressively to subsequent challenges, is crucial for immune system plasticity in disease. Lee et al. [1] recently described a similar kind of immune memory state in astrocytes which they termed “epigenetic memory astrocytes”. The discovery of astrocytes with immune memory could have tremendous importance in understanding and treating neurological disease. However, the RNA-seq data and in vitro experiments presented by Lee et al. to claim astrocytes possess pro-inflammatory immune memory show signs of immune cell contamination. Further, astrocyte-specific knockout of *Ep300*, the purported epigenetic regulator of this memory, did not reduce expression of any memory astrocyte signature genes. The FIND-seq signature used to verify the presence of epigenetic memory astrocytes in experimental autoimmune encephalomyelitis (EAE) also shows signs of immune cell contamination, and the cells identified as memory astrocytes in previously published EAE single-cell RNA-seq data are misannotated macrophages. Lastly, we find the purported epigenetic memory astrocytes identified in single-nucleus RNA-seq data of multiple sclerosis (MS) tissue are an artifact of ambient RNA, low quality nuclei, and non-astrocyte contamination. We conclude that the epigenetic memory astrocyte signature is likely driven by immune cell contamination and the existence of astrocyte immunological memory is insufficiently evidenced. We caution that astrocyte transcriptomic, epigenomic, and functional assays must take care to exclude contamination by immune cells, especially when evaluating the potential of astrocytes to perform immunological functions.

## INTRODUCTION

Astrocytes are multi-functional glial cells of the central nervous system (CNS) with particular relevance to the pathophysiology of neurological disease [2,3]. A hallmark of nearly all insults to the CNS is astrocyte reactivity, a dramatic (yet highly context-dependent) molecular, morphological, and functional remodeling which can result in both loss- and gain-of-function – and can be variably protective or harmful to the health of the CNS [4-6]. Astrocytes are especially potent responders to and effectors of inflammation, responding to microglia- and peripheral immune cell-derived cytokines and secreting a plethora of cytokines and other soluble molecules which can alter neuronal activity and survival [7,8], modulate blood brain barrier permeability [9,10], and impart both pro- and anti-inflammatory effects on microglia and other immune cells [11,12]. Given these prominent roles, astrocytes represent a powerful yet untargeted therapeutic nexus in the management of CNS diseases.

To this end, considerable interest has been directed at understanding the epigenetic mechanisms regulating astrocyte responses to inflammation and other immune stimuli [13-19]. A particularly intriguing question in the field has been whether chronic or repeated exposure to immune stimuli might result in persistent epigenetic changes modifying astrocyte responses [20, 21]. Such a response would be akin to the immune memory of innate immune cells like macrophages (including microglia), monocytes, and dendritic cells, in which a prior immune stimulus “trains” those cells to respond more forcefully to future insults via epigenetic remodeling [22,23]. This memory is a critical component of immune system plasticity and responses to neurological disease [24,25]. Lee et al. recently described an astrocyte state with epigenetically regulated immune memory which they termed “epigenetic memory astrocytes” [1]. Here we reanalyze the presented data to evaluate the evidence for this specialized astrocyte state.

## RESULTS

### Signs of immune cell contamination in cytokine treatment experiments in vivo & in vitro

The description of epigenetic memory astrocytes by Lee et al. is founded primarily on their initial experiment using RNA-seq to profile astrocytes from mice given either one intracerebroventricular (ICV) injection of IL-1β and TNF or two ICV injections one week apart. The authors found an increase in expression of genes consistent with heightened pro-inflammatory responses in astrocytes from the twice challenged mice and identified histone acetyltransferase p300 as a regulator of this response using pathway analysis. With in vitro cytokine treatment assays, the authors concluded that this signature was cell-intrinsic, and thus defined this transcriptional state as “epigenetically controlled memory astrocyte[s]”. From the differentially expressed genes between the twice and once challenged conditions, Lee et al. further defined (see Methods) a signature of genes they termed the epigenetic memory astrocyte “up-signature” and “down-signature”, reflecting genes particularly increased or decreased, respectively, in the twice stimulated condition.

Examining the differential expression test results in Lee et al.’s Supplementary Table 1, we find that the two-hit cytokine stimulus condition displayed higher expression of typical immune cell and myeloid lineage marker genes (e.g. *Cd14, Mpeg1, F13a1, Ms4a4a, Ms4a4b, Tyrobp, Ly6a, Ly6c1, Cl-ec4d, Clec7a, S100a8, S100a9*) and lower expression of canonical astrocyte markers (*Aldh1l1, Slc1a3, Slc1a2, Gfap, Aqp4, Atp1b2, S100b, Sox9, Aldoc, Gja1, Hepacam*) compared to the single stimulus condition (Figure 1a). Examining expression of the up- and down-signature gene lists across the Allen Brain Cell single-cell atlas [26], we found nearly all of the genes comprising the epigenetic memory astrocyte transcriptional signature (the “up-signature”) were most highly expressed in immune cells – monocytes, dendritic cells, border-associated macrophages (BAMs), microglia, and lymphoid cells (Figure 1b-c). In contrast, genes de-enriched in memory astrocytes (the “down-signature”) were mostly highly expressed in astrocytes and astroglial lineage cells like Bergmann glia and astroependymal cells. Using the same gene signature scoring method [27] implemented by Lee et al., we found the up-signature gene set was most highly enriched in immune cell populations (monocytes, dendritic cells, BAMs, microglia, and lymphoid cells) and least enriched in astrocytes, while the down-signature was most enriched in astrocytes and least enriched in immune cells (Figure 1d).

**Figure 1.**
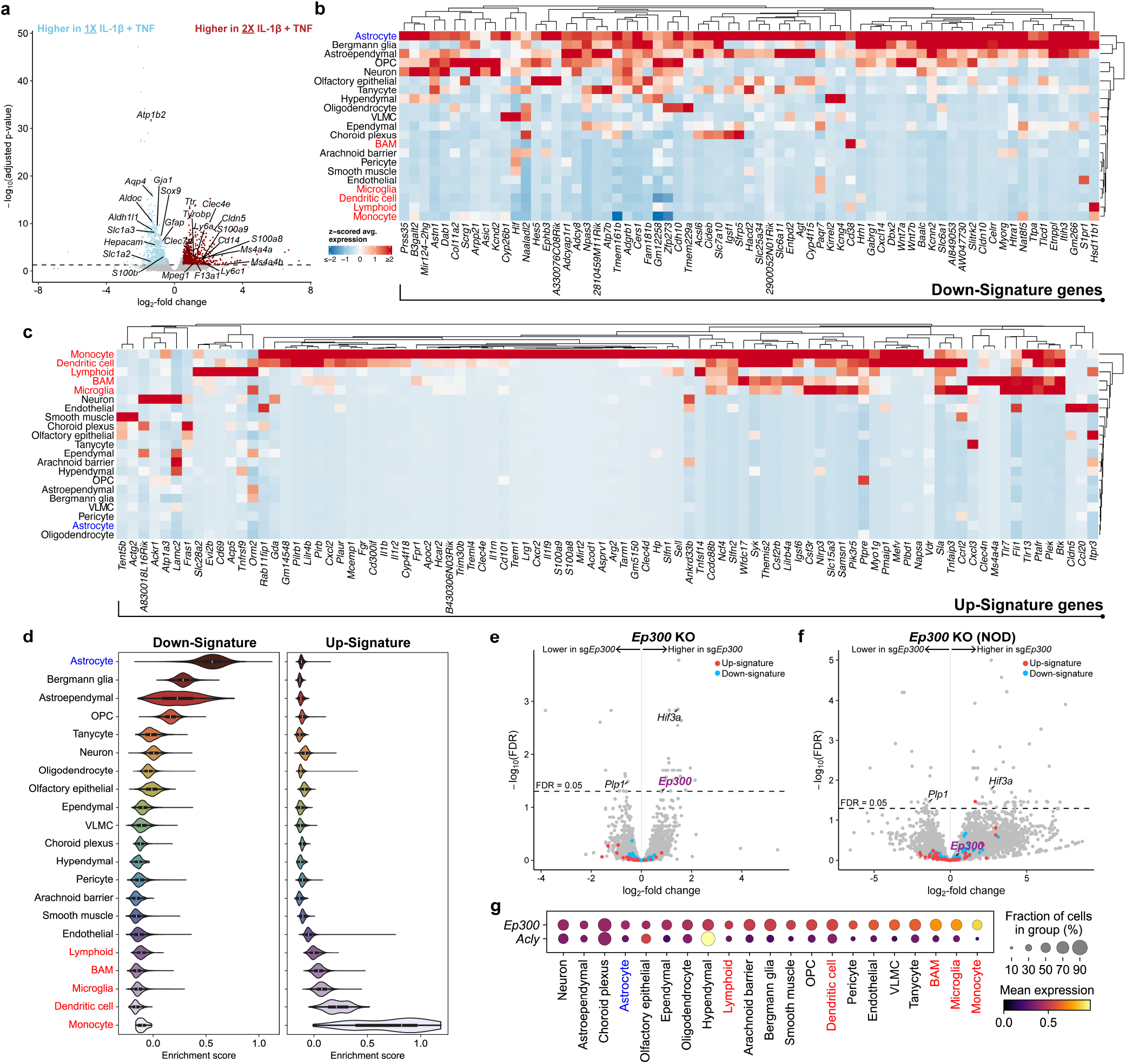
The epigenetic memory astrocyte signature is most concordant with non-astrocyte cell type contamination & p300 knockout did not reduce signature. **(a)** Volcano plot depicting differential expression test results comparing the 2X IL-1β & TNF treated condition and 1X IL-1β & TNF treated condition from Lee et al.’s Supplementary Table 1 (significance threshold shown: adjusted p-value < 0.05, |log2-fold change| > 0.5). Heatmaps of z-scored average expression of the genes comprising the epigenetic memory astrocyte “down-signature” **(b)** and “up-signature” **(c)** defined by Lee et al. across cell type clusters from the Allen Brain Cell Atlas [26]. **(d)** Violin plots of gene set enrichment scores for the up- and down-signatures across the Allen Brain Cell Atlas cell type clusters. **(e)** Volcano plot depicting the differential expression test results comparing sg*Ep300* to sg*Scrmbl* astrocytes. **(f)** Volcano plot depicting the differential expression test results comparing sg*Ep300* to sg*Scrmbl* astrocytes in the NOD EAE experiment. **(g)** Dot plot of expression of *Ep300* and *Acly* across cell type clusters in the Allen Brain Cell Atlas [26].

The Allen dataset is from wild type mouse brain without an inflammatory insult, and it is possible that some of the up-signature genes are restricted to immune cells in the healthy brain but induced in astrocytes during inflammation; however, the majority of these genes have not been demonstrated to be expressed in astrocytes, even in disease. Further, the twice challenged condition demonstrated decreased expression of core astrocyte marker genes like *Gfap, S100b, Sox9*, and *Aldh1l1*, which previous studies have not shown reduced in reactive astrocytes [4, 28, 29]. We believe the simplest explanation for these findings is not that astrocytes are capable of expressing a much larger repertoire of immune genes than previously appreciated; rather, these data likely reflect skewed cell type composition between experimental conditions, perhaps due to the authors’ FACS isolation strategy. Multiple antibodies were used to label and negatively select against non-astrocyte cells, with no positive selection for astrocytes. This strategy assumes that all cells remaining after negative selection are astrocytes, however inefficiencies in antibody labeling inevitably lead to imperfect negative selection, leaving contaminating non-astrocytes in the final isolate. The purity of the primary astrocyte cultures in the in vitro cytokine stimulation experiments, the foundation for the claim that astrocytes possess cell-intrinsic immune memory, is similarly concerning. The authors used a protocol for the culture of primary astrocytes in which mouse brain cortices are dissociated and plated without positive selection of astrocytes or negative selection of non-astrocytes [30]. Removal of non-astrocytes is accomplished by shaking the cultures to detach contaminant cells; however, microglial contamination is an acknowledged limitation of this culture system [30]. The authors performed negative selection gating against CD45+CD11b+ cells (microglia/macrophages) when sorting *Rela*-eGFP+ versus *Rela*-eGFP-cells after the first IL-1β + TNF stimulation in their *Rela*-eGFP experiments (their Extended Data Figure 1m), and this panel clearly shows a notable fraction of cells in the initial cultures were CD45+CD11b+ (up to 15% and 8%, respectively, in the two samples shown).

### Astrocyte-specific knockout of Ep300 did not affect the epigenetic memory signature

Further evidence that this astrocyte state may be artifactual comes from the astrocyte-specific *Ep300* knockout experiments. According to the authors’ Supplementary Table 4, which contains differential expression test results comparing sg*Scrmbl*- and sg*Ep300*-transfected astrocytes, only 10 genes were significantly decreased in expression in the sg*Ep300* group (FDR < 0.05) – and none of these genes were present in the list of “up-signature” genes forming the memory astrocyte signature (Figure 1e). The same is true of the *Ep300* knockout performed in EAE challenged NOD mice. While the volcano plots presented in Lee et al.’s Extended Data Figure 6g depicted the differential expression test results using p-values unadjusted for multiple comparisons (based on the authors’ Supplementary Table 8), only 15 genes were significantly decreased – none of which were components of the up-signature (Figure 1f). Only a single gene was commonly downregulated in the two *Ep300* knockout experiments: the oligodendrocyte-specific myelin component *Plp1* (Figure 1e-f). *Hif3a* was the only commonly upregulated gene, and neither *Hif3a* nor *Plp1* were component genes of the up- or down-signature gene sets. Importantly, *Ep300* expression was unchanged in either of the knockout experiments (Figure 1e-f). Given these issues, there is insufficient evidence that p300 is an upstream regulator of an epigenetic memory astrocyte state.

### Markers used for identifying epigenetic memory astrocytes in situ and by FIND-seq are not specific

The immunostaining data presented indicates an increase in p300 and ACLY, as well as increased H3K27 acetylation, in astrocytes in EAE and multiple sclerosis (MS); however, ACLY and p300 immunoreactivity in EAE and MS tissue is not sufficient evidence of an epigenetic memory state. These factors are expressed across all cell types in the nervous system (including astrocytes) according to the Allen Brain Cell Atlas [26], notably with highest expression in monocytes, microglia, and BAMs (Figure 1g). While the authors used FIND-seq (focused interrogation of cells by nucleic acid detection and sequencing) [14] to demonstrate the concordance of the transcriptional signature of *Ep300*+*Acly*+ astrocytes in EAE with the epigenetic memory astrocyte up-signature, the 661 gene FIND-seq derived signature also shows evidence of immune cell contamination (Extended Data Figure 1a), and it includes several microglia and macrophage-specific genes (e.g. *Csf1r, Cx3cr1, Hexb, Lyz2, Ms4a6c, Tyrobp, Ly6a, Ly6e*, and *Ly86*) and *Ptprc* (CD45) – an exclusive marker of the hematopoietic lineage.

### The epigenetic memory astrocytes identified in EAE scRNA-seq data are macrophages

Concerns about non-astrocyte contamination in the RNA sequencing experiments could be resolved by showing epigenetic memory astrocytes exist using single-cell RNA-seq. In their reanalysis of astrocyte EAE scRNA-seq data from Wheeler et al. [15], Lee et al. identified a cluster (Cluster 3) with increased expression of both the FIND-seq derived and cytokine treatment experiment derived epigenetic memory astrocyte gene signatures compared to the other astrocytes profiled. They further noted that this cluster expanded 18-fold at the peak of EAE. However, the top marker genes for Cluster 3, the population labeled epigenetic memory astrocytes, include macrophage marker genes (*Cd14, Aif1* [IBA1], *Csf1r, Cx3cr1, F13a1, Mpeg1, Ms4a4c, Ms4a6b, Ms4a6c, Ms4a6d*), lymphocyte antigen genes (*Ly6a, Ly6c2, Ly6i, Ly86*), and Fc receptor genes (*Fcer1g, Fcgr1, Fcgr2b, Fcgr3*) (Figure 2a-c). The downregulated marker genes for Cluster 3 include most common astrocyte marker genes (*Slc1a3, Slc1a2, Apq4, Al-dh1l1, Glul, Atp1b2, S100b, Sox9, Aldoc*). It is evident this cluster is not composed of astrocytes but is rather macrophages. The expansion of this cluster at the peak phase of EAE is consistent with the well described expansion of resident macrophages and infiltration of peripheral myeloid cells in EAE [31].

**Figure 2.**
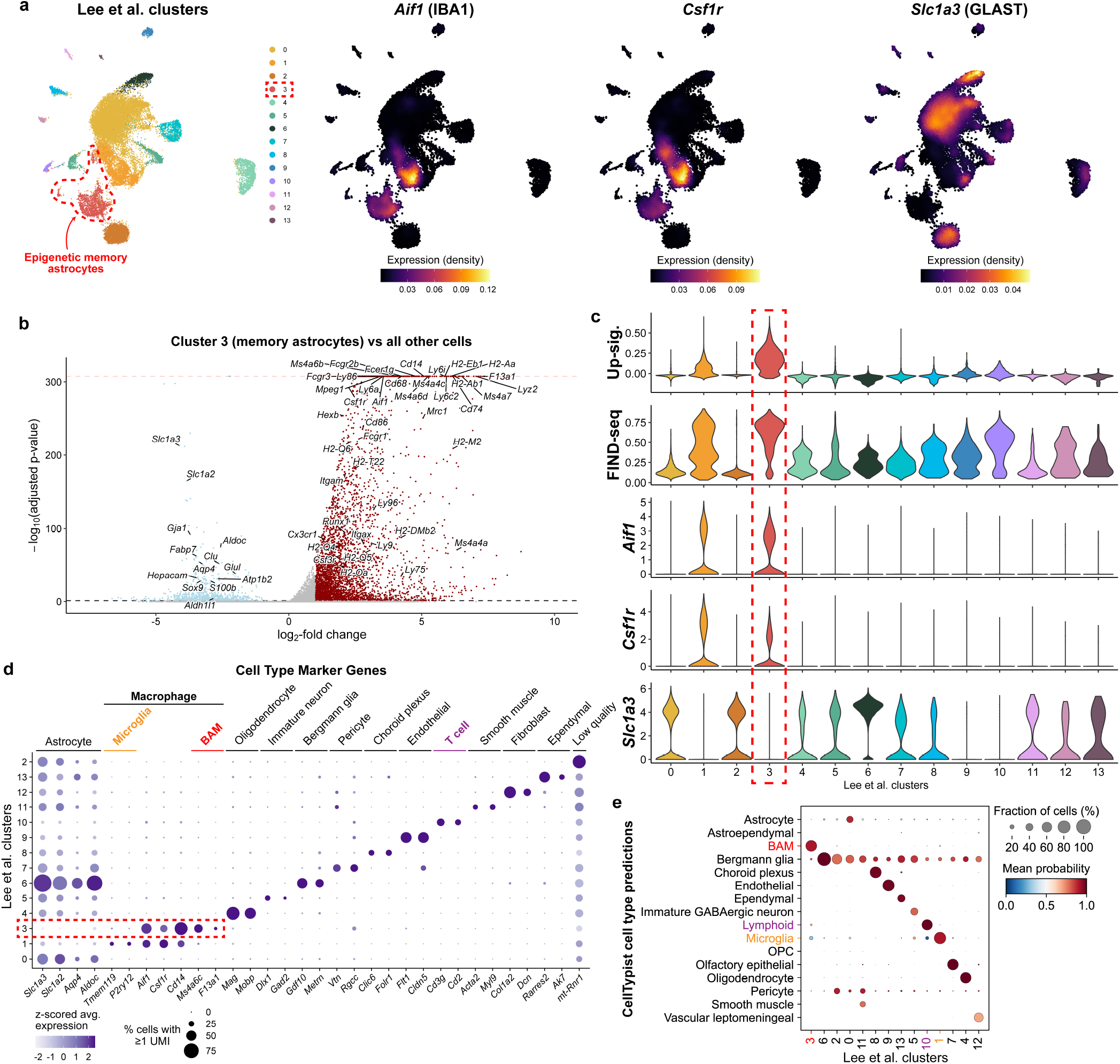
The epigenetic memory astrocytes described in the Wheeler et al. EAE scRNA-seq data are macrophages. **(a)** far left, UMAP embedding depicting clusters reported by Lee et al., which was a reanalysis of previously published data from Wheeler et al. [15]. The memory astrocyte cluster, Cluster 3, is highlighted with a red outline. UMAPs colored by gene expression (depicted by weighted kernel density estimation) of macrophage-specific marker genes *Aif1* (IBA1) (center left) and *Csf1r* (center right) and astrocyte marker gene *Slc1a3* (GLAST) (far right). **(b)** Volcano plot highlighting top positive and negative marker genes for Cluster 3 cells (the cluster labeled as memory astrocytes by Lee et al.) compared to all other cells in the Wheeler et al. dataset. Dotted red line indicates the lowest number representable in the R programming language (i.e. the lowest possible p-value). Note that the x-axis is clipped at ±10 log2-fold change, and a small number of genes with fold changes exceeding ±10 are thus not shown. See our Supplementary Table 1 for complete list of differentially expressed genes. **(c)** Violin plots depicting expression of *Aif1, Csf1r*, and *Slc1a3*, as well as enrichment scores for the up-signature and FIND-seq derived gene signature across each cluster. The purported epigenetic memory astrocyte cluster is outlined in red. **(d)** Dot plot of canonical cell type marker genes across the clusters. The purported epigenetic memory astrocyte cluster is outlined in red. Cluster 2 was identified as low quality cells on the basis of highly elevated proportions of UMIs mapping to the mitochondrial genome. **(e)** Dot plot illustrating the automated cell type annotations assigned to the cells in each of the Lee et al. clusters by CellTypist [33].

Mis-annotation of non-astrocyte cell types in this dataset was not limited to Cluster 3. The authors referred to Clusters 1 and 10 as “pathogenic astrocyte subsets expanded in EAE”; however, Cluster 1 was marked by several of the same pan-myeloid genes expressed in Cluster 3 (*Cd14, Aif1, Csf1r, Cx3cr1*), as well as microglia-specific [32] genes *Tmem119* and *P2ry12* (Figure 2d, Supplementary Table 1). These cells are microglia. This cluster includes nearly all of the cells originally defined by Wheeler et al. as a distinct MAFG-driven astrocyte subtype present in EAE and MS (Extended Data Figure 1b-c; labeled Cluster 5 in [15]). The cells in Cluster 10 are T cells, delineated by T cell-specific marker genes like *Cd2, Cd3g, Cd4, Trbc1, Trbc2*, and *Trac* (Figure 2d, Supplementary Table 1). The expansion of Clusters 1 and 10 thus reflects the known expansion of these immune cell populations in EAE [31]. Most other clusters in this dataset also appear to be non-astrocyte cell types based on canonical marker gene expression (Figure 2d, Supplementary Table 1). In support of our manual annotations, we found the automated cell type annotation tool CellTypist [33] labeled Cluster 3 as BAMs, Cluster 1 as microglia, and Cluster 10 as lymphoid cells rather than astrocytes (Figure 2e). Given the extensive cell type mis-annotation, these data do not support the existence of an epigenetic memory astrocyte population in EAE.

### The epigenetic memory astrocyte signature identified in MS snRNA-seq data is an artifact

Lastly, the authors combined two prior single-nucleus RNA-seq (snRNA-seq) datasets generated from postmortem tissue from patients with multiple sclerosis or healthy controls [34, 35] and evaluated expression of the epigenetic memory astrocyte gene signature in these datasets. They discovered Cluster 2 astrocytes most highly expressed the FIND-seq derived and cytokine treatment experiment derived memory astrocyte gene signatures and found this cluster was most prevalent in chronic lesion samples from MS patients. However, examining the top marker genes for each cluster, we found several clusters were mis-annotated non-astrocytes (Figure 3a-b). Cluster 3 appeared to correspond to ependymal cells (*CFAP299, TTC6, SPAG17*), Cluster 4 to oligodendrocytes (*MBP, MOBP, MYRF*), and Cluster 5 to neurons (*SYT1, SNAP25, SLC17A7*). When examining Cluster 2’s top marker genes, we found that while this cluster was marked by increased expression of *GFAP* and *VIM*, it was also defined by markers of lymphocytes, particularly T cells (*THEMIS, FYB1, SKAP1, IKZF1*) and B cells (*IGHGP, IGKC, IGHG1, IGHG3, IGLC2*) (Figure 3c). While Cluster 2 exhibited highest expression of the FIND-seq gene signature (Figure 3d), we found that the up-signature enrichment scores for this cluster were actually lower than Cluster 3 (ependymal cells) (Figure 3e, Supplementary Table 1), and nearly all of the up-signature genes were not expressed in Cluster 2 or any of the other clusters in the dataset (Extended Data Figure 2a).

**Figure 3.**
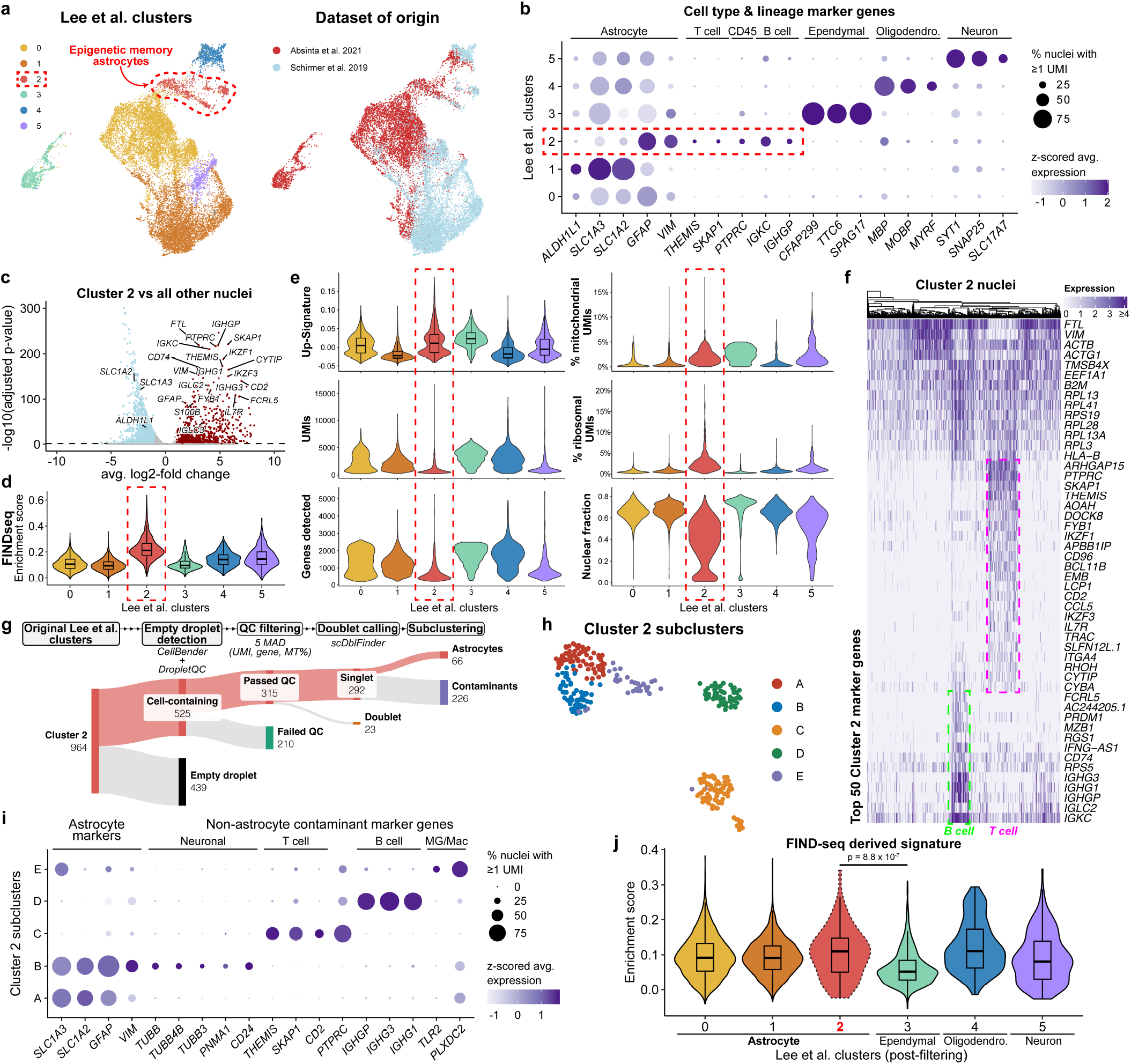
The epigenetic memory astrocytes described in MS snRNA-seq data are composed of empty droplets, low quality outliers, and non-astrocyte cell types. **(a)** left, UMAP embedding depicting clusters reported by Lee et al. reanalyzing published data from Absinta et al. [34] and Schirmer et al. [35]. The cluster labeled as epigenetic memory astrocytes (Cluster 2) is highlighted with a red outline. right, UMAP embedding colored by dataset of origin. **(b)** Dot plot depicting expression of select cluster marker genes, with cell type/lineage specific marker genes annotated. **(c)** Volcano plot of Cluster 2 marker genes compared to all other nuclei in the merged dataset. See our Supplementary Table 1 for complete list of differentially expressed genes. **(d)** Violin plot displaying FIND-seq gene signature enrichment scores across clusters. **(e)** Violin plots depicting the up-signature enrichment scores (see Supplementary Table 1 for statistics), number of UMIs, number of genes with expression detected, percentage of UMIs mapping to the mitochondrial genome, as well as ribosomal genes, and the proportion of intronic UMIs (nuclear fraction) per nucleus across clusters. **(f)** Heatmap depicting normalized expression of the top 50 positive marker genes (ranked by adjusted p-value) for Cluster 2 across all nuclei in the cluster. Columns are clustered based on Euclidean distance in principal component analysis (PCA) space. **(g)** Sankey schematic depicting the proportion of Cluster 2 which were excluded following re-analysis by CellBender and DropletQC empty droplet calling, >5 MAD quality control filtering, scDblFinder doublet calling, and the removal of contaminant non-astrocyte subclusters. **(h)** UMAP embedding depicting results of sub-clustering the remaining Cluster 2 nuclei. **(i)** Dot plot of expression of canonical cell type marker genes across Cluster 2 sub-clusters. **(j)** Violin plot showing FIND-seq derived epigenetic memory astrocyte signature enrichment scores across all clusters following QC filtering and Cluster 2 contaminant removal. P-value corresponds to Wilcoxon rank-sum tests where Cluster 2 was compared pairwise against all other clusters, with Bonferroni correction for multiple comparisons. All comparisons not shown were non-significant (p > 0.05), see our Supplementary Table 1 for full results.

We also noted that Cluster 2 appeared to exhibit signs of low quality or damaged nuclei, including decreased UMIs (unique molecular identifiers, i.e. transcripts) and genes detected per nucleus, increased proportions of reads mapping to the mitochondrial genome and ribosomal genes, and decreased nuclear fractions (the proportion of reads mapping to introns rather than exons) (Figure 3e). By examining the top 50 markers for this cluster in a heatmap (Figure 3f), we noticed many of these genes, particularly the T cell and B cell markers, were expressed by only a subset of Cluster 2 cells. This suggested Cluster 2 included contaminant non-astrocyte cell types but might not be entirely comprised of non-astrocytes, requiring further analysis.

To evaluate whether Cluster 2 contained any bona fide astrocytes expressing epigenetic memory signature genes, we first performed ambient RNA correction and empty droplet classification, sample-level outlier removal, and doublet/multiplet removal (see Methods). We found that 69.6% of nuclei from Cluster 2 were excluded after applying these standard QC metrics, leaving just 292 remaining nuclei (Figure 3g, Extended Data Figure 2b). We next performed sub-clustering, identifying five distinct populations – three of which were delineated by T cell (*THEMIS, SKAP1, CD2*), B cell (*IGHGP, IGHG3, IGHG1*), and microglia/macrophage (*TLR2, PLX-DC2*) genes, respectively (Figure 3h-i, Extended Data Figure 2c, Supplementary Table 1). Two remaining subclusters exhibited the highest expression of astrocyte genes (subclusters A & B); however, subcluster B also expressed markers of neurons and neural progenitors (including *PNMA1, CD24, TUBB, TUBB3*, and *TUBB4B*). We thus classified subcluster A as bona fide astrocytes and subclusters B through E as contaminants. When the contaminant subclusters were removed, we found the remaining Cluster 2 contained only 66 nuclei (6.8% of the original cluster), with 10 or fewer nuclei per sample (Extended Data Figure 2d). Further, Cluster 2 did not exhibit significantly elevated enrichment of the epigenetic memory astrocyte gene signatures compared to other astrocyte or non-astrocyte contaminant clusters (Figure 3j, Extended Data Figure 2e-f, Supplementary Table 1). We conclude that the elevated expression of the epigenetic memory astrocyte gene signatures in these human MS datasets reported by Lee et al. is an artifact driven by ambient RNA contamination, low quality or damaged nuclei, and contamination by non-astrocyte cell types. We thus believe the presented evidence does not support the existence of epigenetic memory astrocytes in multiple sclerosis.

## DISCUSSION

The identification of an astrocyte state with epigenetically regulated immune memory could profoundly impact the understanding and treatment of diseases like MS; however, the data presented by Lee et al. is insufficient to establish the existence of such a state. We found signs of immune cell contamination in the RNA-seq data which were the basis for this claim and the definition of a distinct gene signature. We saw evidence of immune cell contamination in the in vitro experiments foundational to the claim that such memory is cell-intrinsic and epigenetically regulated. We found insufficient evidence for the claim that knockout of *Ep300* in astrocytes suppressed the epigenetic memory gene signature. And we identified errors in cell type annotation and insufficient quality control in the sc/snRNA-seq analysis used to claim an epigenetic memory astrocyte state is present in EAE and MS. We thus conclude the epigenetic memory astrocytes described by Lee et al. are likely an artifact of immune cell contamination.

These issues highlight the challenges of studying astrocytes using transcriptomic, epigenomic, and functional assays. While astrocytes have numerous roles in the healthy and diseased nervous system, studies of these roles have been complicated by difficulties in isolating pure and viable populations of astrocytes for ex vivo transcriptomic profiling or in vitro assays [36]. Methods for generating primary astrocyte cultures for in vitro studies have historically resulted in non-astrocyte cell type contamination or the induction of immature and reactive transcriptional states in astrocytes [37, 38]. Over the past decade, isolation of highly pure populations of astrocytes from mouse brain has become increasingly feasible due to the creation of well-validated astrocyte-specific antibodies [39, 40] and astrocyte-labeling fluorescent reporter and TRAP mouse lines (e.g. MGI:3843271; MGI:J:150403) [41-43], which can be used for positive selection [28, 29, 44, 45]. Similarly, the development of a primary astrocyte isolation and culture method using immunopanning greatly reduced the occurrence of non-astrocyte contamination and immature or reactive gene expression in in vitro experiments [38].

Lee et al. opted to use a negative selection strategy to deplete non-astrocyte cells rather than positive selection when isolating astrocytes for transcriptomic and epigenomic profiling. We believe studies profiling astrocytes ex vivo should use well-established positive selection tools whenever possible to minimize non-astrocyte cell type contamination. When this is not possible, non-astrocyte contamination should be explicitly assessed. Lee et al. also opted to use a protocol which results in microglia contamination [30] when generating primary astrocyte cultures. Given the variability in purity between different methods of culturing astrocytes, we believe studies using astrocytes in vitro should include data assessing the purity of the cultures used.

Many of our concerns could have been allayed if the authors had unambiguously demonstrated expression of their epigenetic memory signatures in astrocytes with single-cell RNA-seq. However, the cell populations identified by the authors as epigenetic memory astrocytes in the reanalyzed single-cell datasets were almost entirely composed of misannotated non-astrocyte cell types and empty or low quality droplets – highlighting the need for more explicit standards for quality control and cell type annotation in single-cell analysis. Further, the authors failed to show clear expression of their memory astrocyte gene signatures in these data. The single-cell gene signature scoring method used by Lee et al., originally described by Tirosh et al. [27], derives its enrichment score by simply averaging the expression of genes in the signature and subtracting from this value the average expression of a larger set of control genes within each single cell. With this approach, gene signature enrichment scores can be elevated by expression of just a single gene, giving the appearance of enrichment even when nearly all of the signature genes are not expressed – exemplified by Extended Data Figure 2a & e-f.

To avoid mistaken conclusions, we believe studies defining and applying gene signatures across transcriptomics datasets ought to demonstrate robust expression of a significant fraction of the proposed signature’s genes before asserting its presence in a different dataset. Ideally, this should include evaluating whether the individual genes of the signature are differentially expressed in the examined context with an appropriate statistical test. For single-cell RNA-seq experiments, this should include pseudobulk differential expression tests or mixed models which account for variation between biological replicates and minimize false discoveries due to pseudoreplication [46, 47].

While the evidence presented by Lee et al. is insufficient to establish the existence of an epigenetic memory astrocyte state, we do not mean to claim that astrocytes are incapable of forming an immune memory-like response to disease stimuli. Future studies should continue to explore this possibility, with careful attention to the danger of immune cell contamination in astrocyte isolations and primary cultures.

## SUPPLEMENTARY INFORMATION

**Supplementary Table 1**. Excel spreadsheet containing **(a)** Wilcoxon rank-sum test results of differential expression between each cluster defined by Lee et al. and all other cells in the Wheeler et al. dataset; **(b)** Wilcoxon rank-sum test results of differential expression between each cluster defined by Lee et al. and all other nuclei in the merged Absinta et al. and Schirmer et al. dataset; **(c)** Wilcoxon rank-sum test results comparing up-signature enrichment scores between Cluster 2 and each other cluster defined by Lee et al. and all other nuclei in the merged Absinta et al. and Schirmer et al. dataset. **(d)** Wilcoxon rank-sum test results of differential expression between each Cluster 2 subcluster and all other nuclei from our filtered reanalysis of the human snRNA-seq data; **(e)** Wilcoxon rank-sum test results comparing FIND-seq signature enrichment scores between Cluster 2 and each other cluster after empty droplet removal, QC filtering, doublet removal, and contaminant removal; **(f)** Wilcoxon rank-sum test results comparing up-signature enrichment scores between Cluster 2 and each other cluster after empty droplet removal, QC filtering, doublet removal, and contaminant removal.

## METHODS

No original data were collected in the preparation of this manuscript. Original analyses were conducted with R (version 4.3.3) and Python (version 3.12.8) and are described below.

### Comparing the epigenetic memory astrocyte signature to the Allen Brain Cell Atlas

#### (Figure 1a-d, Figure 1g, Extended Data Figure 1a)

We obtained metadata and H5AD files containing processed data from the Allen Brain Cell Atlas corresponding to the 10X-v2, 10X-v3, and 10X-multi dataset subsets using the abc_atlas_access package (v3.1.2, release 11-30-2024) [26]. Original cluster “class” labels were aggregated into broad cell type groupings (e.g. merging all neuronal clusters into one “Neuron” category). The “up-signature” and “down-signature” gene sets were derived based on the methods and code described in correspondence with the authors, who stated that their Supplementary Table 3 (originally described in the paper as containing the up- and down-signatures) included incorrect genes. Instead, genes increased or decreased in expression (|log2-fold change| > 1.5, FDR < 0.05) in the 2X IL-1β + TNF versus 1X IL-1β + TNF conditions from the in vivo cytokine treatment RNA-seq experiment which were also increased or decreased, respectively, (|log2-fold change| > 1, FDR < 0.05) in the 2X IL-1β + TNF versus 2X PBS control condition formed the up-signature and down-signature, respectively. The 2X IL-1β + TNF versus 1X IL-1β + TNF differential expression test results were obtained from Lee et al.’s Supplementary Table 1, and the 2X IL-1β + TNF versus 2X PBS differential expression test results were obtained in correspondence with the authors. Enrichment scores across cell type categories in the Allen Brain Cell Atlas data were calculated using the Scanpy [48] function score_genes, which applies the enrichment scoring method originally described by Tirosh et al. [27] which is also implemented in the Seurat [49] function AddModuleScore, which was used by Lee et al. for scoring enrichment of gene set signatures across single cells/nuclei from the sc/snRNA-seq datasets. For Extended Data Figure 1a, enrichment scores were calculated using identical methods for the 661 gene FIND-seq signature from Lee et al.’s Supplementary Table 9. Violin plots in Figure 1 and Extended Data Figure 1a were produced with Scanpy (v1.10.4) [48]. Figure 1 heatmaps were produced with ComplexHeatmap (v2.18.0) [50].

### Reviewing the Ep300 knockout RNA-seq results

#### (Figure 1e-f)

Volcano plots were generated from the differential expression test results in Lee et al.’s Supplementary Table 4 for the *Ep300* knockout experiment using ggplot2. We permissively defined significantly up- and down-regulated genes in the sg*Ep300* condition compared to the *Scrmbl* control as those with a multiple comparisons-adjusted p-value < 0.05 with no minimum log2-fold change threshold. Identical methods were used for examining the NOD EAE *Ep300* knockout RNA-seq data from Lee et al.’s Supplementary Table 8.

### Reanalysis of the mouse EAE scRNA-seq data

#### (Figure 2, Extended Data Figure 1b-c)

Reanalysis of the mouse EAE scRNA-seq data from Wheeler et al. [15] was performed with Seurat (v5.1.0) [49] using a preprocessed Seurat object provided to us by Lee et al., which contained the cluster labels and UMAP and t-SNE dimension reductions from the original manuscript. UMAP plots overlaid with gene expression depicted by weighted kernel density estimation were produced with the Nebulosa package (v1.12.0) [51]. Differential expression testing (Wilcoxon rank-sum test) between each cluster and all other cells was performed using the wilcoxauc function from the presto R package (v.1.0.0) [52], and log2-fold changes were calculated using the formula used by the Scanpy package, a choice made based on recent work discussing the differences between fold change calculations between the Seurat and Scanpy packages [53]. The Cluster 3 differential expression volcano plot was produced with ggplot2. The x-axis was clipped at ± 10 log2-fold change to make visualization of most data points easier; however, we note this resulted in the exclusion of a small number of data points from the plot. See our Supplementary Table 1 for complete list of differentially expressed genes. Violin plots and dot plots were produced with Seurat. Automated cell type annotation was performed with CellTypist (v1.6.3) [33] using a reference model created from the Allen Brain Cell Atlas dataset [26]. Cell type categories with fewer than 400 cells (representing fewer than 0.01% of the cells in the 4 million cell dataset) were excluded from the reference, and the remaining cell type groups were randomly downsampled such that each cell type was represented by no more than 5,000 single cells. The CellTypist “train” function was run on the downsampled dataset using default parameters. Raw count data from the Wheeler et al. dataset Seurat object provided by Lee et al. was used as input for annotation with the resulting model, with predictions made by majority voting. The Sankey plot in Extended Data Figure 1c was created from a table provided by the original authors containing, for each cell barcode, the cluster identities presented in the Wheeler et al. [15] and Lee et al. papers and was produced using SankeyMATIC (https://sankeymatic.com).

### Reanalysis of human MS snRNA-seq data

#### (Figure 3, Extended Data Figure 2)

Initial reanalysis of the human MS snRNA-seq data (originally from Absinta et al. [34] and Schirmer et al. [35]) was performed with Seurat using a preprocessed Seurat object provided to us by Lee et al. Enrichment scores for the up-signature and FIND-seq derived gene signature were calculated with the AddModuleScore function, with the original mouse gene sets converted to one-to-one human orthologs using the orthogene package (v1.8.0) [54]. We note the enrichment scores we calculated for each single cell may differ from those presented by Lee et al., as the authors attempted to convert from mouse genes to human orthologs simply by capitalizing the mouse gene names. This approach results in inaccurate conversion of some mouse gene names, thus resulting in different enrichment scores due to loss of the mis-converted genes. Statistical significance for gene signature enrichment between clusters was tested using Wilcoxon rank-sum tests for pairwise comparison of Cluster 2 with all other clusters, with p-values corrected for multiple comparisons using the Bonferroni correction method (with results shown in Supplementary Table 1). Differential expression testing (Wilcoxon rank-sum test) between each cluster and all other nuclei was performed using the wilcoxauc function from the presto package, and log2-fold changes were calculated using the formula used by the Scanpy package, as described for the mouse analysis above. The Cluster 2 differential expression volcano plot was produced with ggplot2, and the x-axis was clipped at ± 10 log2-fold change to make visualization of most data points easier; however, we again note this resulted in the exclusion of a small number of data points from the plot. See our Supplementary Table 1 for complete list of differentially expressed genes. Normalized expression of the top 50 marker genes (by adjusted p-value) for all Cluster 2 nuclei was plotted in a heatmap using ComplexHeatmap, with the columns (nuclei) ordered according to clustering based on Euclidean distances in Harmony [55] batch-corrected PCA space. Rows were ordered using the seriation R package (v1.5.7) [56]. Dot plots of expression of cluster markers or signature genes were produced with the Seurat DotPlot function.

The original FASTQ files for the Absinta (GSE180759) and Schirmer (PRJNA544731) datasets were obtained from the Sequence Read Archive (SRA) using the fasterq-dump function from the sra-tools package (v2.11.0). Both datasets were processed using Cell Ranger (v7.0.1, 10X Genomics). Ambient RNA removal and empty droplet classification were performed with CellBender (v0.3.0) [57], with a false positive rate (FPR) of 0.05 and learning rate of 5 × 10^−6^. Additional empty droplet detection was performed using the identify_ empty_drops function from the DropletQC package (v1.0.0) [58] using the Cell Ranger produced BAM files for each sample as input. Nuclear fractions for all barcodes were calculated using the DropletQC function nuclear_fraction_tags. All droplets labeled as empty by either CellBender or DropletQC were classified as empty and excluded from further analysis. Code from the scCustomize R package (v3.0.1) [59] was adapted to create a Seurat object from the CellBender output files.

The isOutlier function from the scater package (v1.30.1) [60] was used to remove nuclei which exceeded 5 median absolute deviations (MADs) from the within-sample median for number of UMIs, number of genes detected, and percentage of UMIs mapping to the mitochondrial genome for each nucleus. This threshold has been recommended by a recent “best practices” review of single-cell analysis methods and has been described as a “lenient” quality control (QC) threshold [61, 62]. For the Absinta et al. dataset, a single sample exhibited mitochondrial UMI percentages far higher than all other samples, and these outlier nuclei were not classified as such by the 5 MAD threshold due to a high within-sample median. Nuclei in this sample were thus subjected to an additional manual threshold, removing any nuclei with a mitochondrial UMI percentage greater than the maximum mitochondrial UMI percentage among all other samples after the 5 MAD thresholding. Nuclei classified as outliers along any of the above QC metrics were excluded from further analysis. scDblFinder (v1.16.0) [63] was used to identify doublets and multiplets among the remaining nuclei, and downstream analysis was performed only with nuclei labeled as singlets. The Sankey plot in Figure 3g was produced using Sankey-MATIC. Importantly, all of the above preprocessing steps were applied to the complete Absinta et al. and Schirmer et al. datasets, not just the cells originally contained in Lee et al.’s analysis.

The remaining Cluster 2 cells were subsetted for further analysis. The top 2000 variable features were calculated using the FindVariableFeatures Seurat function with selection method “dispersion”. The expression data was then rescaled, PCA was performed, and the Harmony R package (v1.2.3) was used to correct for batch differences across tissue sample and dataset of origin. A nearest neighbors graph was created using the FindNeighbors function using the first 10 Harmony-corrected principal components, and sub-clustering was performed using the FindClusters function with a resolution parameter of 0.5. Five clusters (labeled with alphabetical identifiers “A” through “E”) were found using this method, and cluster marker genes were calculated by Wilcoxon rank-sum test as described above. Based on the Wilcoxon test results, only subcluster A exhibited expression of astrocyte marker genes without expression of non-astrocyte marker genes. Subcluster A was retained as bona fide astrocytes while all other subclusters were excluded as non-astrocyte contaminants.

After removal of contaminants from Cluster 2, the Add-ModuleScore function was again used to calculate enrichment scores for the up-signature and FIND-seq gene signatures across the final clusters. Statistical significance was tested using Wilcoxon rank-sum tests for pairwise comparison of Cluster 2 with all other clusters, with p-values corrected for multiple comparisons using the Bonferroni correction method.

## DATA & CODE AVAILABILITY

No original data were collected for the analyses in this manuscript. Data in this manuscript include supplementary tables originally published in Lee et al., data from the Allen Brain Cell single-cell atlas [26] (available from Amazon Web Services at https://allen-brain-cell-atlas.s3.us-west-2.amazonaws.com/index.html), differential expression test result tables and preprocessed single-cell Seurat data objects provided to us by the original authors of Lee et al., and original single-cell RNA-seq data from Wheeler et al. [15] (available from the NCBI GEO repository at accession GSE129763), Absinta et al. [34] (available at NCBI accession GSE180759), and Schirmer et al. [35] (available at NCBI accession PRJNA544731). All code necessary for the retrieval of these input files and reproduction of the analyses and figure panels in this manuscript is available in a Github repository in the form of annotated Jupyter notebooks: https://github.com/michael-r-odea/EpiMemAstros. We have made all input files not retrieved from AWS or NCBI available in a Zenodo repository: https://zenodo.org/records/14919413.

## AUTHOR CONTRIBUTIONS

M.R.O. performed the data analysis. M.R.O. and S.A.L. wrote the manuscript.

## ACKNOWLEDGEMENTS

We thank Itai Yanai and James Salzer (NYU Grossman School of Medicine), Philip Hasel (University of Edinburgh), and anonymous members of the community for their insightful advice and review of this manuscript. We thank Christopher Glass (UC San Diego) for general advice on currently accepted practices for generation and analysis of epigenetic data. M.R.O. was supported by a predoctoral fellowship from the National Institute of Neurological Disorders and Stroke of the National Institutes of Health (F31NS135948).

---

**Extended Data Figure 1.**
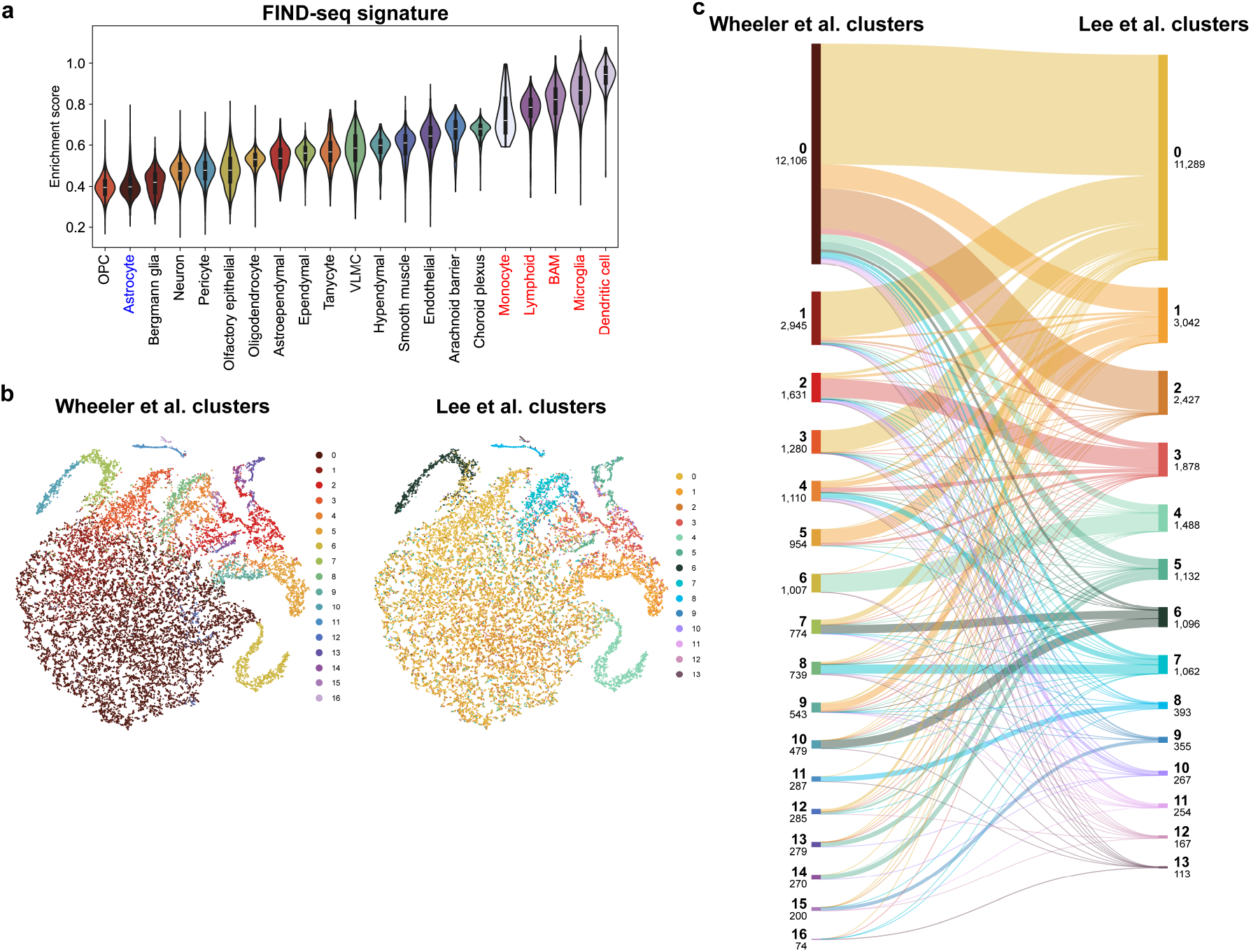
**(a)** Violin plot of gene set enrichment scores for the FIND-seq derived epigenetic memory astrocyte gene signature across the Allen Brain Cell Atlas cell type clusters. **(b)** t-SNE embedding from Wheeler et al. [15], with cells colored by Wheeler et al. clusters (left) or Lee et al. clusters (right) for the mouse EAE scRNA-seq data. **(c)** Sankey diagram demonstrating the overlap between the clusters originally defined in Wheeler et al. [15] and clusters from the reanalysis of Lee et al.

**Extended Data Figure 2.**
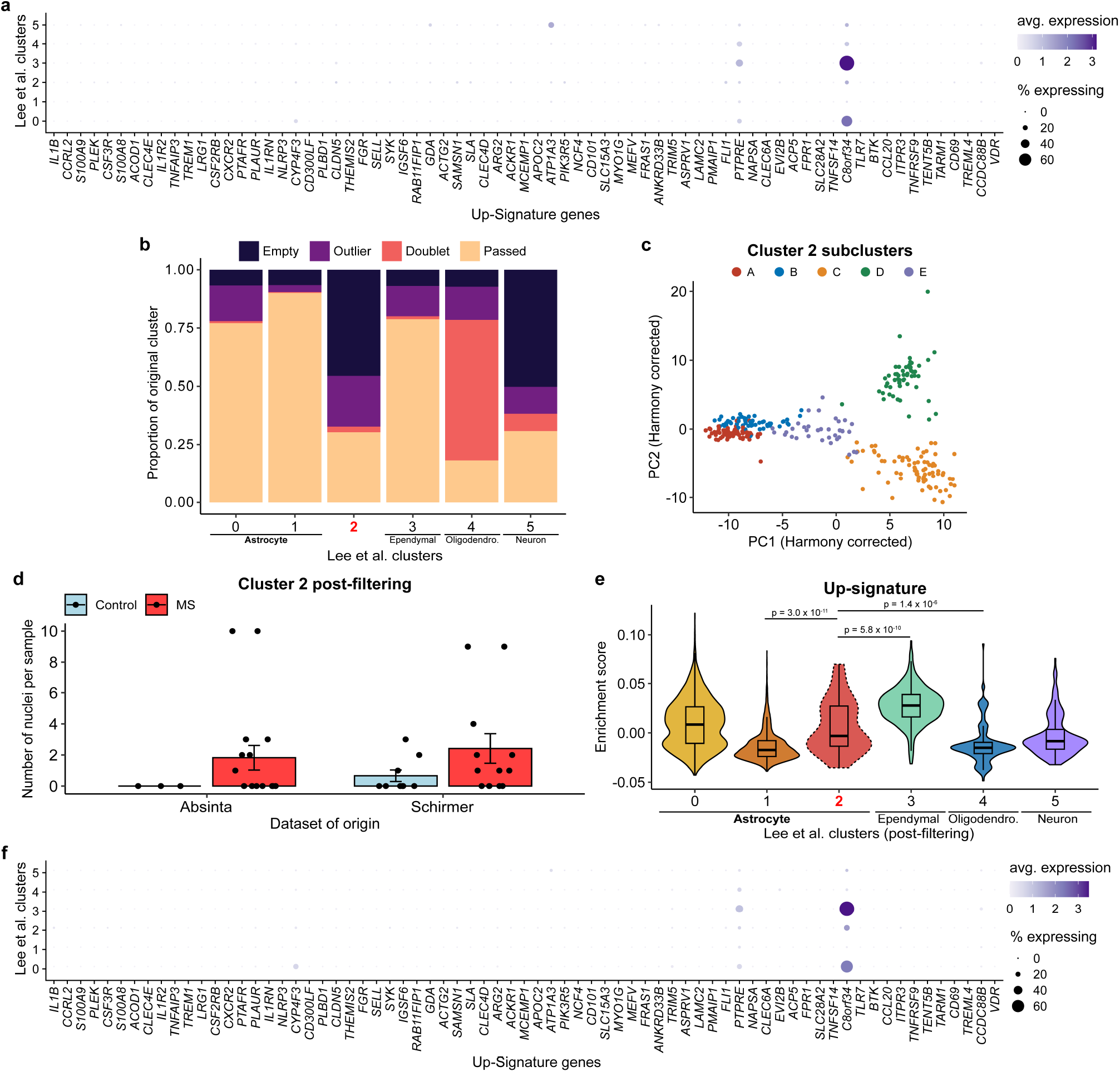
**(a)** Dot plot showing expression of all one-to-one orthologous genes from the up-signature gene set across the clusters reported by Lee et al. with the human single nucleus RNA-seq data from Absinta et al. [34] and Schirmer et al. [35]. **(b)** Bar plot depicting the proportion of nuclei from each cluster which were attributed to empty droplets by CellBender and DropletQC (“Empty”), which failed the 5 median absolute deviations (MAD) quality control thresholding (“Outlier”), or which were called by scDblFinder as doublets or multiplets (“Doublet”). The remaining nuclei which passed all three preprocessing steps are labeled “Passed”. **(c)** Scatter plot of Cluster 2 subclusters in Harmony-corrected PCA space. **(d)** Quantification of the number of nuclei from each tissue sample in the final filtered Cluster 2, split across dataset and donor diagnosis. Error bars depict standard error of the mean. **(e)** Violin plot of gene set enrichment scores for the cytokine stimulation derived epigenetic memory astrocyte up-signature across the clusters reported by Lee et al. after applying empty droplet removal, QC filtering, doublet removal, and removal of non-astrocyte contaminants from Cluster 2. P-value corresponds to Wilcoxon rank-sum tests where Cluster 2 was compared pairwise against all other clusters, with Bonferroni correction for multiple comparisons. All comparisons not shown were non-significant (p > 0.05), see our Supplementary Table 1 for full results. **(f)** Dot plot showing expression of all one-to-one orthologous genes from the up-signature across clusters following empty droplet and ambient RNA removal, QC filtering, doublet removal, and removal of non-astrocyte contaminants (the latter performed only for Cluster 2).

